# A chloroplast cell-free system for measuring ribosome binding site strengths

**DOI:** 10.1101/2024.02.17.580832

**Authors:** Lauren Clark, Christopher A. Voigt, Michael C. Jewett

## Abstract

Plastid engineering offers the potential to carry multi-gene traits in plants, however, it requires reliable genetic parts to balance expression. The difficulty of chloroplast transformation and slow plant growth make it challenging to build plants just to characterize genetic parts. To address these limitations, we developed a cell-free system from *Nicotiana tabacum* chloroplast extracts for prototyping genetic parts. Our cell-free system uses combined transcription and translation driven by T7 RNA polymerase and works with plasmid or linear template DNA. To develop our system, we optimized lysis, extract preparation procedures (e.g., runoff reaction, centrifugation, and dialysis), and the physiochemical reaction conditions. Our cell-free system can synthesize 34 ± 1 μg/mL luciferase in batch reactions. We apply our system to test a library of 104 ribosome binding site (RBS) variants and rank them based on cell-free gene expression. We observe a 1300-fold range of luciferase expression normalized by mRNA expression, as assessed by the malachite green aptamer (relative luminescence units per relative fluorescence units). We also find a positive correlation between the observed expression in chloroplast extracts and the predictions made by the RBS calculator. We anticipate that chloroplast cell-free systems will increase the speed and reliability of building genetic programs in plant chloroplasts for diverse applications.

## Introduction

Plant chloroplasts represent a maturing frontier of synthetic biology, with opportunities to produce pharmaceuticals, nutrients, and biosensors. Plant synthetic biology has been slower to develop than its bacterial counterpart.^1^ Plant genetic engineering is much more difficult and lengthier than bacteria (e.g., chloroplast transformation can take six to twelve months to achieve homoplasty and then seeds^2^). In addition, many species are not amenable to genetic manipulation and have limited sets of genetic tools.^3,4^ This backdrop has resulted in a paradigm where useful genetic designs of plant origin are often ported into more tractable organisms such as *Escherichia coli* or *Chlamydomonas* for evaluation before going through the trouble of moving the pathway to a plant. However, non-plant systems differ in their molecular composition and regulatory signals, which can lead to inaccuracies in the data generated.

Cell-free gene expression (CFE) systems^5,6^ have recently proven useful for accelerating biological design in the context of their native host organism’s biological machinery.^7–15^ In recent years, for example, the development of CFE systems derived from a diverse set of organisms such as *Pseudomonas*,^16^ *Streptomyces*,^17–21^ *Vibrio natriegens*,^22,23^ *Saccharomyces*,^24^ *Clostridium*,^25^ *Pichia pastoris*,^26^ and Chinese Hamster Ovary^27,28^ cells have opened new opportunities in rapid prototyping of synthetic biological systems, including genetic parts, such as promoters and ribosome binding sites.^15,16,29^

Here, we set out to advance CFEs for high-throughput genetic part prototyping in plant chloroplasts. We focused on chloroplasts for several reasons. First, they are not subject to generational silencing, which can be a problem with transformation into the plant nucleus. Second, chloroplasts are known for producing high titers of recombinant protein.^4,30–34^ Third, chloroplast expression offers an inherent biocontainment^35,36^ because chloroplasts are inherited maternally.^37^ Finally, plastids have their own genome and bacteria-like ribosomes, which make them well-suited for cell-free expression.

As a model, we developed an optimized CFE platform from purified *Nicotiana tabacum* chloroplasts. This builds off decades of work in chloroplast molecular biology that have been a primary means of elucidating chloroplast genetics.^38–40^ Previous works, for example, have shown the ability to purify chloroplasts from spinach and tobacco,^41–45^ and demonstrated the preparation of translation-only capable chloroplast extracts,^46,47^ as well as *in vitro* transcription and translation.^41,42^ These early *in vitro* systems proved useful for the analysis of promoter binding and transcription, leading to a greater understanding of chloroplast promoter architecture and polymerase-DNA binding.^38,39,48,49^ Additionally, chloroplast extracts^50^ and extract-based expression systems have been used to elucidate chloroplast ribosome binding and understand the biology of translation initiation in the chloroplast.^46,51–53^

We established a protocol for producing highly active chloroplast extracts, optimized the chemical reaction environment to increase CFE yields of reporter proteins, and then applied the method to screen a library of ribosome binding sites (RBSs) designed using the RBS calculator.^54–59^ We discovered that chloroplast extracts maintain mRNA over long periods of time (greater than 24 h), which we predict could be a useful feature for applications in testing circuits and sensors.

## Results and Discussion

### Activating high yielding protein production in cell-free chloroplast extracts

The goal of this work was to develop CFE system from *N. tabacum* chloroplasts capable of manufacturing easy-to-use reporter proteins for genetic part prototyping. We grew *N. tabacum* plants in Conviron growth chambers with a 16 h-8 h light-dark cycle at 28 °C.^47^ Tobacco leaves were harvested after 6 weeks, and chloroplasts were purified from the three youngest, fully-expanded leaves using a blender, cloth straining, and density centrifugation (**Figure 1A**). After purification, chloroplasts were then flash frozen and stored at -80°C. Then, chloroplasts were lysed, and extracts were processed based on previous protocols for translation-only extracts.^46,47^ These extracts had 32 ± 5 mg/mL of total chloroplast protein and were subsequently tested for activity in cell-free transcription and translation.

**Figure 1.**
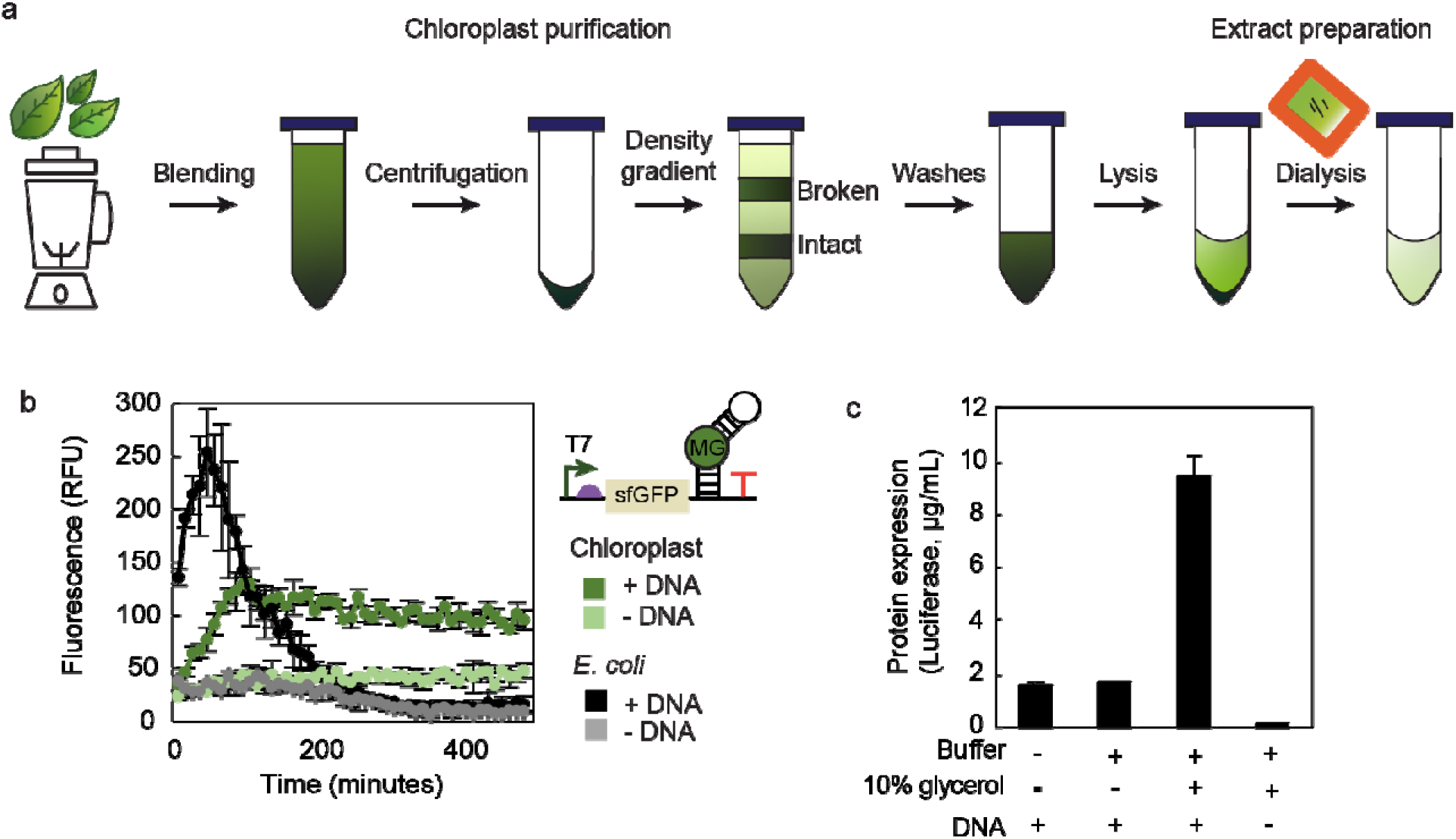
Establishing a cell free transcription and translation system from *Nicotiana tabacum* chloroplasts. (A) Schematic of chloroplast purification and extract preparation. (B) Cell-free transcription of the malachite green RNA aptamer (MGA) from a T7 RNAP promoter in a CFE reaction containing the malachite green dye using either chloroplast or *E. coli* lysates and with or without plasmid DNA. (C) Combined cell-free transcription and translation reactions from chloroplast extracts are active. A no DNA negative control CFE reaction is shown (-DNA). Values show means with error bars representing standard deviations (s.d.) of at least three independent experiments.

We first carried out cell-free transcription in a reaction mixture containing a DNA template, exogenously added T7 RNA polymerase (RNAP), energy substrates, nucleotides, and salts necessary for gene expression.^47,60^ We assessed transcription levels with a reporter template that harbored the malachite green aptamer (MGA) sequence (**Figure 1B**). Once transcribed, this RNA aptamer binds to malachite green and activates the dye’s fluorescence.^61,62^ mRNA concentrations remain stable for over 20 h in chloroplast extracts. This contrasts to *E. coli* extracts, where RNA increases rapidly, peaks, and degrades (**Figure 1B**), as has been observed before.^15,63^

We then assessed combined transcription and translation of a luciferase reporter protein. Luciferase was selected because bioluminescence assays are highly sensitive and could be useful for genetic part prototyping. Unfortunately, our early protein expression levels were low (∽2 μg/mL) (**Figure 1C**). In making extracts for those reactions, we directly froze the chloroplast pellets after purification but before lysis. We worried that chloroplasts might break open after thawing but prior to extract preparation (**Figure 1A**), reducing the translation machinery available for protein production in prepared extracts. Therefore, we next resuspended chloroplasts in lysis buffer or lysis buffer with 10% glycerol lysis prior to freezing in liquid nitrogen and extract preparation (**Figure 1C**). Glycerol can act as a cryoprotectant to protect chloroplasts from freezing damage and premature rupture. We found that resuspending chloroplasts in a 10% glycerol lysis buffer yielded extracts with 30% higher total protein concentrations (42 ± 2 mg/mL), and improved CFE productivity 5-fold. Protein expression yields could be further improved by adjusting codon usage of our pJL1 luciferase reporter,^64^ which was originally created for bacterial expression, to be optimized for chloroplast expression (**Supplementary Figure 1**). We used this optimized coding sequence and chloroplasts prepared in glycerol for all further cell-free reactions.

### Optimizing the cell-free reaction environment

We next sought to increase protein synthesis yields by systematically optimizing the physiochemical reaction environment (**Figure 2A**). This was important because the physiochemical conditions of cell-free reactions are known to play a key role in the operation of *in vitro* biological processes.^65–67^

**Figure 2.**
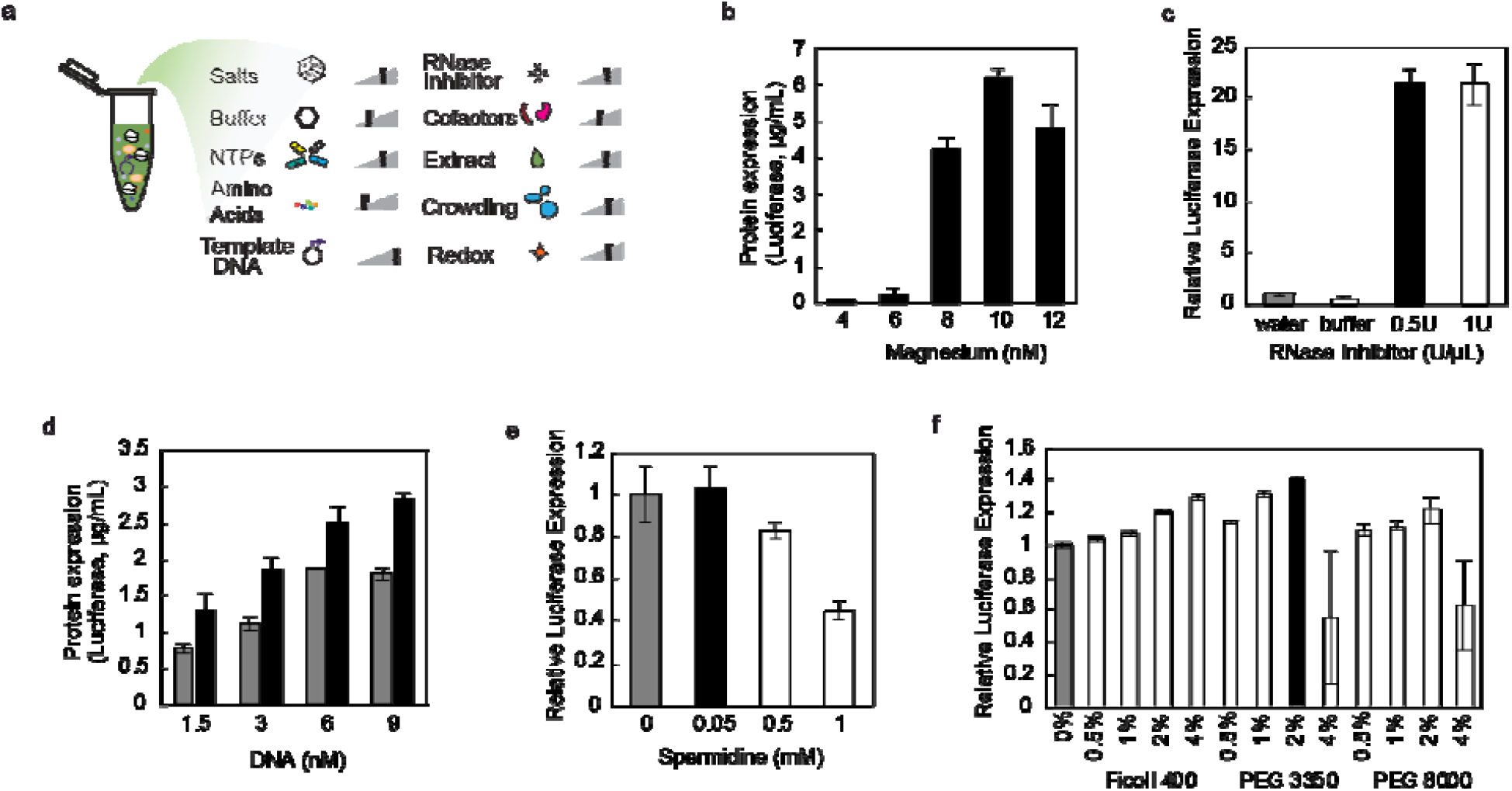
Physiochemical optimization. (A) The cell-free environment was optimized by testing a range in concentrations of several reaction components. These components include: (B) magnesium, (C) RNase inhibitor, (D) DNA in the form of plasmid (gray bars) or linear expression templates (LET, black bars), (E) spermidine, and (F) crowding agents. Notably, magnesium is optimized as a quality control step for each independent extract. Gray bars indicate the previous condition prior to optimization and black bars indicate the condition used in all future experiments. Values show means with error bars representing standard deviations (s.d.) of at least three independent experiments.

Our initial physiochemical optimizations focused on optimizing the salt concentrations, buffer, extract amount, and reduction potential. We started with salts and specifically magnesium, which has been previously shown to be a critical component of CFE reactions.^65^ Our data suggested an optimum magnesium concentration of 10 mM (**Figure 2B**). However, we later found that re-optimization of magnesium was needed for each batch of extract, which is typical in the field.^27,65,68,69^ After magnesium, we optimized potassium, which is used as the major cation of the system, as well as ammonium, which is used to mimic the natural environment. We found that 60 mM potassium acetate and 30 mM ammonium acetate led to the highest CFE yields (**Supplementary Figure 2**). Next, we explored a range of buffer concentrations (HEPES 15-120 mM). Our results indicated that the 15 mM HEPES at pH 7.3 led to the highest translational activity. Finally, we assessed the impact of percent extract volume and the reducing agent DTT, settling on 50% volume fraction and 5 mM DTT (**Supplementary Figure 3**).

**Figure 3.**
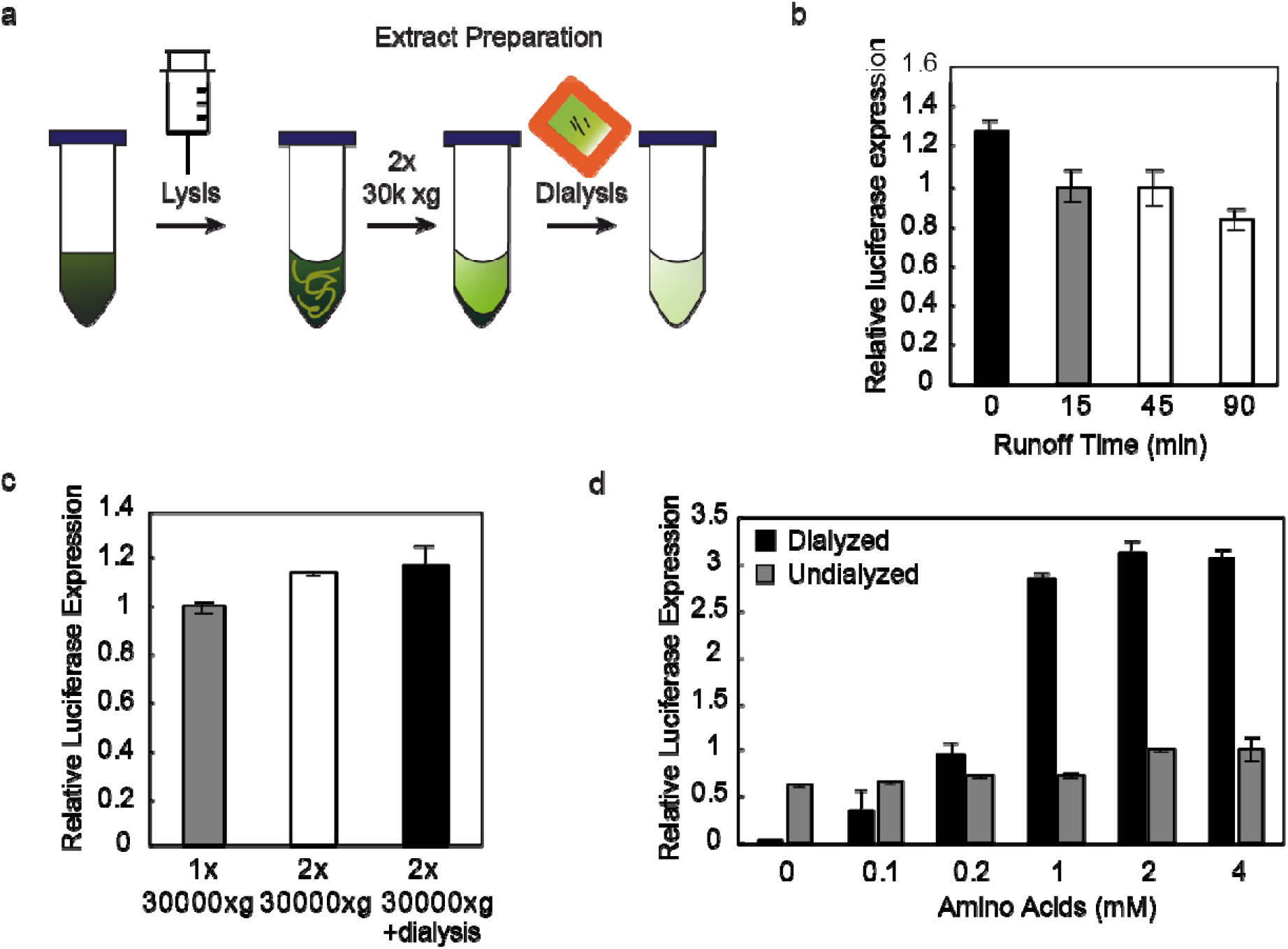
Optimization of extract preparation procedures and amino acids levels. (A) Cartoon schematic of the finalized extract preparation procedure. Our initial extract preparation was modified to remove the runoff incubation, include a second 30000 x g spin, and dialysis. To improve cell-free protein synthesis yields, we optimized extract preparation procedures including: (B) runoff reaction, (C) centrifugation and dialysis, and (D) amino acids. Values show means with error bars representing standard deviations (s.d.) of at least three independent experiments.

We next explored components involved in supporting high-level transcription and translation. We focused on protein coding templates in the form of DNA and RNA. First, we tested adding RNAse inhibitor to our CFE reactions. We observed a dramatic improvement in cell-free activity with 0.5 U/μL RNAse inhibitor (**Figure 2C**). Second, by titrating different concentrations of plasmid DNA, we observed that we could increase yields with maximum expression occurring for plasmid DNA at concentrations of 6 to 9nM (**Figure 2D**). Third, given robust expression with plasmid DNA, we next tried linear DNA expression templates (LETs), in other words PCR products. Relative to plasmid DNA, PCR products have the benefit of avoiding cloning steps and thus can expedite high-throughput workflows for validating genetic part performance *in vitro*. We found that we could increase luciferase yields by over 50% than when plasmid DNA was used as a template (**Figure 2D**). Fourth, we assessed the impact of spermidine on the system, as this polycation has been shown to stabilize DNA, RNA, and tRNA and aid in T7 RNAP function.^70–72^ While spermidine did not statistically increase yields (**Figure 2E**), we elected to add 0.05 mM in subsequent reactions to better mimic cellular physiochemical conditions.^65^ Finally, we assayed a panel of crowding agents given that these impart molecular crowding effects that can enhance transcription and translation activity.^61^ We tested a range of concentrations of Ficoll 400, PEG3350, and PEG 8000 from 0.5% to 4% (v/v) and saw the biggest improvement from PEG 3350 at 2% (**Figure 2F**).

Taken together, our optimization efforts led to an increase in CFE activity of more than 5-fold. **Supplementary Table 1** shows this comparison and the final optimized chloroplast CFE.

### Optimizing cell-free extract production

With new physiochemical conditions at hand, we set out to optimize extract preparation procedures to further increase cell-free protein biosynthesis yields. Our initial procedure consisted of chloroplast lysis, a runoff incubation, and centrifugation at 30,000 x *g* (**Figure 1A**). However, these conditions were not optimized. Here, we examined these extract preparation conditions, including the addition of a dialysis step, starting with the runoff reaction (**Figure 3A**).

The runoff reaction was selected for optimization first because it is hypothesized to release actively translating ribosomes from the thylakoid membrane. We theorized that these might be released in a runoff reaction, improving the translational capacity of our extracts. Given that runoff reactions are thought to allow for translation of endogenous mRNAs to complete and free up available ribosomes for use in cell-free reactions, we reasoned that we might increase available ribosomes by testing different lengths of runoff reactions. We found that omitting any incubation post-lysis was the most beneficial condition (**Figure 3B**).

After removing the runoff reaction step, we found that the chloroplast membrane material pelleted loosely in the glycerol buffer. Thus, we assessed the incorporation of a second centrifugation step at 30,000 x *g* to eliminate membrane material more easily during extract preparation. We observed a slight increase in protein expression yields (**Figure 3C**), but more importantly more robust extract preparation with this step and so it was subsequently always used.

We next explored the impact of dialysis, which is commonly used in CFE protocols to remove metabolic byproducts and provide a suitable storage buffer.^61^ Dialysis was carried out at 4ºC in 10% glycerol lysis buffer (**Figure 3C**). Following dialysis, we conducted a final study to optimize amino acids for the chloroplast CFE reaction. We observed that amino acid concentrations higher than 1 mM were important for achieving high yields in dialyzed extracts (**Figure 3D**).

In sum, by optimizing chloroplast lysis conditions, the physiochemical reaction environment, and extract processing methods, we improved protein synthesis yields more than 100-fold relative to our first starting conditions (**Figure 4A**). **Figure 4B** shows active luciferase yield throughout the duration of the cell-free batch reaction. The final yield of luciferase after a 24-hour incubation in a batch CFE reaction was 34 ± 1 μg/mL. We then applied semicontinuous cell-free protein synthesis reaction^73^ using a dialysis device (3.5 K MWCO) to increase luciferase production yield. Semicontinuous reactions allow for any toxic byproducts to diffuse away from the reaction and any cofactors that are consumed are replenished due to the presence of a semipermeable membrane between the reaction and a buffer that includes a subset of the reaction components. With a semicontinuous setup, luciferase production was increased to 60 ± 4 μg/mL, which was nearly double the yield of batch reaction conditions (**Figure 4B**). The time-course reactions show that this chloroplast system can produce stable, active protein with little apparent degradation in the case of the enzyme luciferase.

**Figure 4.**
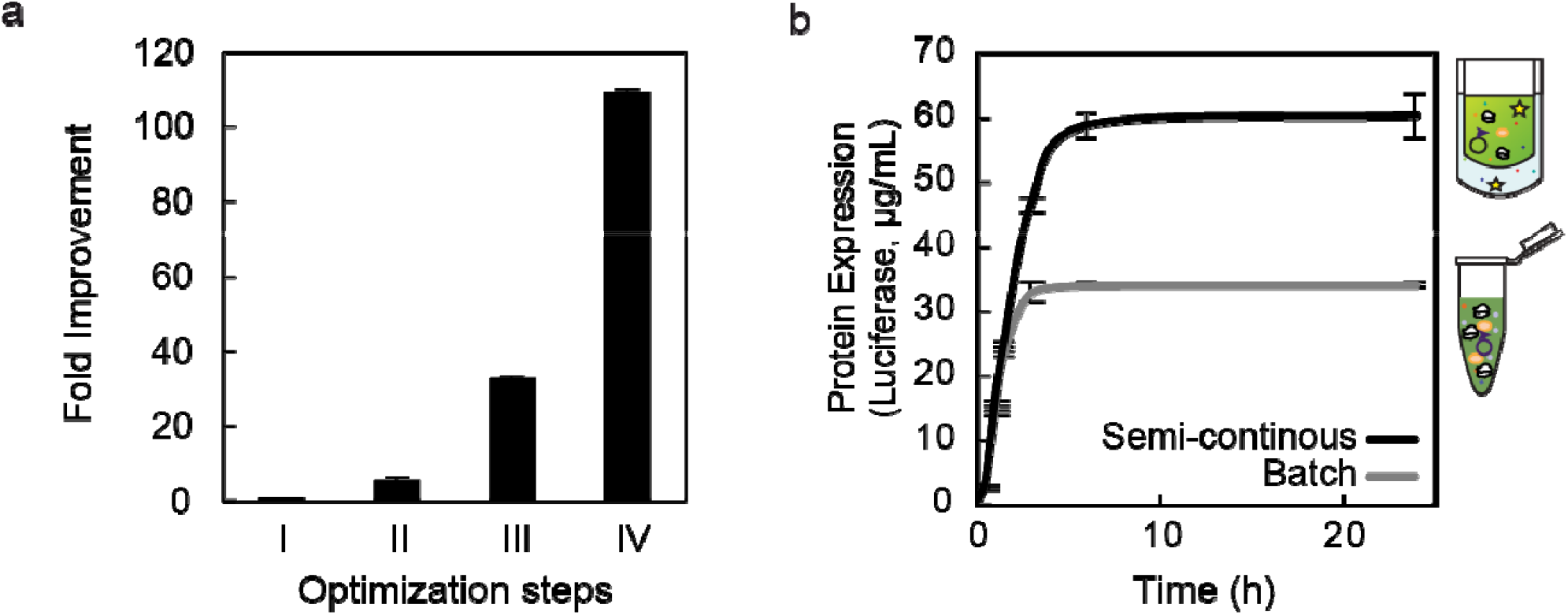
Analysis of improvements and maximum cell-free expression yields. (A) Cumulative improvements of cell-free expression over process (II, glycerol lysis; IV, process optimization) and physiochemical (III) optimizations. (B) Reaction dynamics of batch and semicontinuous reaction modes. Values show means with error bars representing standard deviations (s.d.) of at least three independent experiments.

### Analysis of an RBS library with the chloroplast CFE

We measured the strengths of an RBS library to understand how the cell-free system would perform in a high-throughput analysis of genetic parts. To carry out this test, we designed 104 RBSs replacing the original RBS sequence between the T7 promoter sequence and the start of the luciferase gene (**Figure 5A**). The MGA followed the luciferase gene to allow mRNA measurement. The RBS designs were predicted by the Ribosome Binding Site Calculator^54–59^ with a wide spectrum of expression levels (**Supplementary Figure 4**) using the predicted sequence of AAGGAGVBHDHYBD for the RBS and spacer region. This sequence was sufficiently variable to generate thousands of sequences, while still containing the canonical core sequence of the tobacco chloroplast 16S rRNA.^74^ The resulting library covered a three orders of magnitude range in predicted translation initiation rates (TIRs) from maximal (RBS sequence 1) to minimal (RBS sequence 104) (**Supplementary Table 2**).

**Figure 5.**
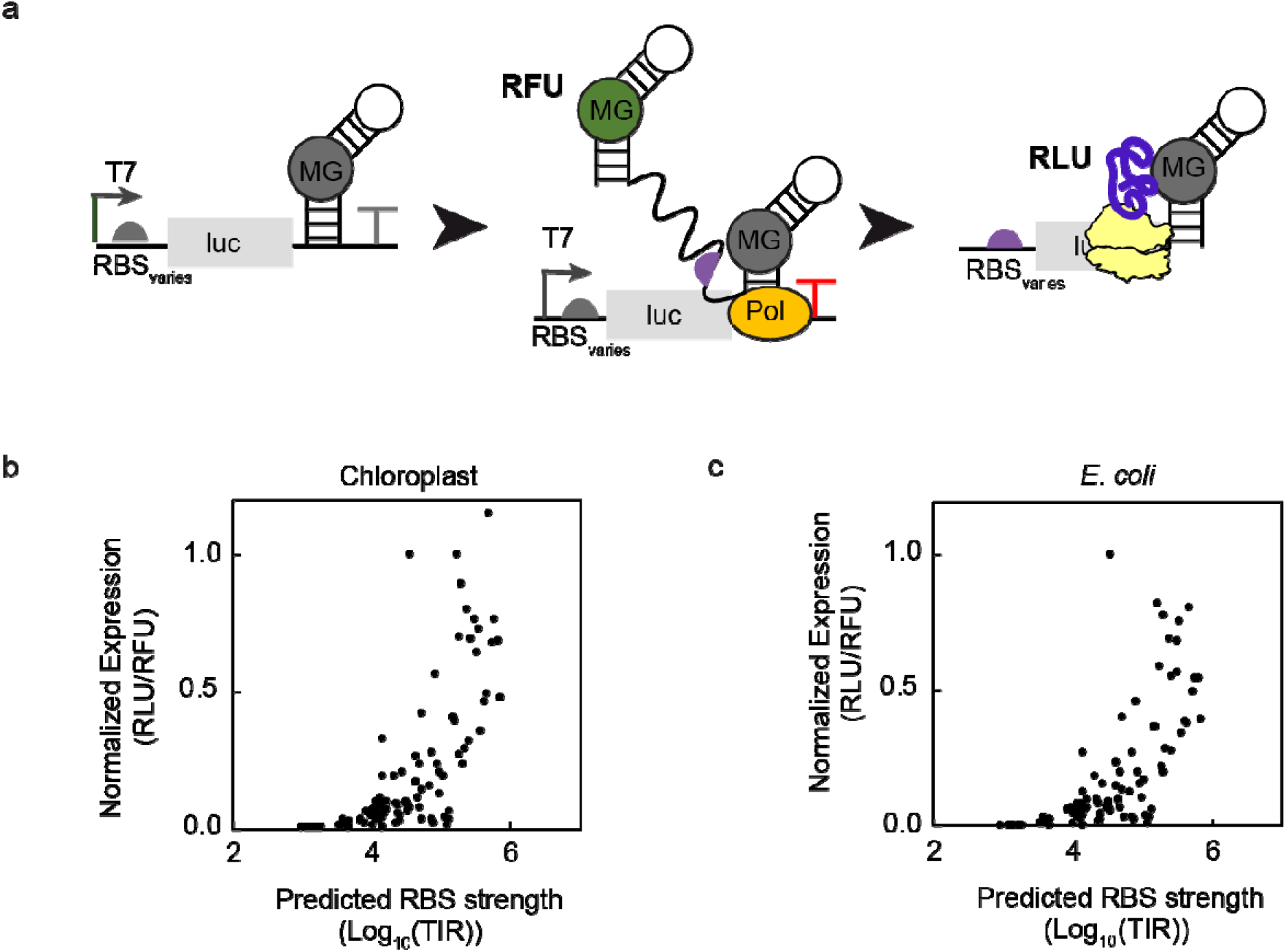
The chloroplast cell-free system can be used to rank genetic parts. (A) Cartoon schematic of the RBS library assay. We compared predicted RBS strength from the Salis calculator versus normalized gene expression, or proteins per transcripts (maximum luminescence/maximum fluorescence), in chloroplast (B) and *E. coli* (C) cell-free systems. Values show means of at least three independent experiments (n=3) with normalized error reported in the supplement.

We carried out cell-free gene expression reactions and assessed both transcription (MGA) and translation (luciferase) activity (**Figure 5**). We observed a 1300-fold range of normalized luciferase expression (proteins/transcripts; relative luminescence units (RLU)/relative fluorescence units (RFU)). We also found a positive correlation between the predicted RBS TIR and luciferase yield normalized by mRNA transcript levels (**Figure 5B**). We observed a correlation between the RBS calculator predicted translation initiation rate (TIR) and normalized cell-free gene expression (R^2^=0.50), noting that poorly performing predicted RBSs matched experimental expression levels well. For comparison purposes, reactions were carried out also in *Escherichia coli* BL21 Star (DE3) extracts (**Figure 5C**). We found similar trends between chloroplast CFE expression and *E. coli* CFE expression (**Supplementary Figure 5**), likely because chloroplast ribosomes contain a prokaryotic-like Shine-Dalgarno (SD) sequence located upstream of the initiator AUG.^50^

## Discussion

We developed a cell-free gene expression system from *Nicotiana tabacum* chloroplasts capable of producing the easy-to-use reporter protein luciferase. Transcription was obtained by the exogenous T7 RNAP. Obtaining sufficient reporter levels was achieved by optimizing plant growth and lysis, the physiochemical reaction conditions, and extract preparation procedures (e.g., runoff reaction, centrifugation, and dialysis). A key insight was the use of glycerol to stabilize the chloroplasts prior to lysis. Batch protein biosynthesis yields of luciferase for the best performing CFE system were 34 ± 1 μg/mL.

The CFE system was applied to screen a library of 104 RBSs in less than one day that were computationally designed using the RBS calculator. We observed a correlation between the RBS calculator predictions of TIR and normalized cell-free gene expression when adjusted for transcription (R^2^=0.50), which was comparable to what we observed with *E. coli* extracts (R^2^=0.47). The library we produced showed a 1300-fold dynamic range of proteins per transcripts between the lowest expressing and the highest expressing sequences, demonstrating a system well suited to ranking DNA templates. The differences found between the predicted expression and the actual expression could be due to several reasons. For example, chloroplast translation varies based on whether the plant is exposed to light or dark, and our workflow was developed using light-harvested chloroplasts. Modifications to the system are needed for the study of nighttime translational programs. Additionally, regulation in plastids occurs at the transcript level by RNAses, so it is not currently known how much of this machinery is present or active in these extracts. Of note, chloroplast RBSs have frequently been tested in living *E. coli*. Our data (**Supplementary Figure 5**) suggest that *E. coli* CFE serves a good model for chloroplast CFE expression.

While we attempted to do so, we did not observe endogenous transcriptional activity in the chloroplast CFE system (data not shown). Previous CFE efforts in *E. coli* have shown that cell-free expression of genes under native bacterial σ70 promoters is constrained by the rate of transcription.^61^ Looking ahead, chloroplast extract preparation procedures could be further modified to explore these phenomena and develop strategies to activate endogenous transcription.

In sum, our work provides a platform for prototyping plant-based genetic parts in a chloroplast CFE system before evaluating smaller design sets in cells, as has been done in a variety of cell free systems.^14,15,75–77^ We anticipate the chloroplast CFE system will accelerate the characterization of reliable genetic parts for plant synthetic biology.

## Methods

### Growth Conditions

Nicotiana tabacum var. Bright Yellow (PI 552597) were acquired from GRIN-GLOBAL and grown on Metro-Mix 360 from SunGro via Fosters, Inc. (Waterloo, IA, USA) at 28°C under 16h white light/8h dark conditions in a Conviron A1000 growth chamber for 6 weeks post-germination.

### Purification of Chloroplasts

Chloroplast purification was adapted from previous work.^47^ The top 3 healthy leaves below the apical leaf were removed from 6-week-old plants exposed to 6-7 h light. 300 g leaves were collected and blended in 100g batches. Leaves were removed to 4 °C for the remainder of the protocol. Leaves were ripped into 4-6 pieces each, and loaded into a Waring blender. 300mL buffer MCB1 (50mM HEPES/KOH, pH 8.0, 0.3 M mannitol, 2mM EDTA, 5mM β-mercaptoethanol) with 0.1% w/v BSA and 0.6% polyvinylpyrrolidone (average molecular weight 40,000) was poured over the leaves, and the blender was run on high for two 5-s intervals, then a 2-s interval, checking the blending at each pause to ensure all leaves are destroyed. The brei was then combined and strained with two sheets each of cheesecloth and Miracloth (EMD Millipore, Burlington, MA, USA) and centrifuged at 1000xg. After centrifugation, the pellet was resuspended in 18 mL of MCB1 with 0.1% BSA and 2-4.5 mL of material was layered onto stepwise Percoll (GE Healthcare, Chicago, IL, USA) MCB1 gradients with 0.1% BSA. Per 300 g leaves, 10 gradients were prepared with a Hamilton syringe with 7 mL of 20% Percoll, 12 mL of 50% Percoll, and 11 mL of 80% Percoll. Loaded Percoll gradients were centrifuged in a fixed-angle rotor for 10 minutes with minimum acceleration and deceleration and the green band between the 50% and 80% phases was collected as intact chloroplasts. Chloroplasts were washed three times in MCB2 (50 mM HEPES/KOH, pH 8.0, 0.32 M mannitol, 2 mM EDTA, 5 mM β-mercaptoethanol). The first wash was in 3:1 volumes of buffer to chloroplast material, the second wash was in 60 mL, and the final wash was in 8mL. To collect chloroplasts between washes, chloroplasts were centrifuged at 1,000 x g for 4 minutes, and after the final wash, they were centrifuged at 5,000 x g for 4 minutes. Between washes resuspension was done by gentle swirling motion by hand to avoid lysing the chloroplasts. After the final wash, chloroplasts were resuspended by gentle pipetting in 1 mL/g lysis buffer (30 mM HEPES/KOH, pH 7.7, 60 mM potassium acetate, 7 mM magnesium acetate, 60 mM ammonium acetate, 10% v/v glycerol, 5 mM DTT, 20 μM each of 20 amino acids, 0.1 mM GTP, and 0.5 mM PMSF, where the amino acids and GTP are added after thawing) and flash frozen, then stored at -80 °C.

### S30 Preparation of Extract

All procedures were carried out at 4 °C or on ice. Unless otherwise noted, frozen chloroplasts in lysis buffer were thawed on ice for 20 minutes, then mixed by pipetting. Chloroplasts were lysed by passing through a 25G syringe 12 times and centrifuged at 30,000 x g at 4 °C for 30 minutes. The supernatant was removed and centrifuged a second time at 30,000 x g at 4 °C for 30 minutes. The supernatant from the second spin was loaded into a dialysis cassette (Slide-A-Lyzer 10K MWCO, Pierce Biotechnologies, Waltham, MA, USA), and dialyzed twice for 2 h each against 600 mL of buffer with 30 mM HEPES/KOH, pH7.7, 60 mM potassium acetate, 7 mM magnesium acetate, 60 mM ammonium acetate, 10% v/v glycerol, 5 mM DTT, and 0.5 mM PMSF, and centrifuged a final time at 4 °C at 30,000 x g for 20 minutes. The supernatant was removed and aliquoted, then flash frozen and stored at -80 °C.

### Cell-Free Protein Synthesis Reaction

Reactions were run at 25 °C in 10 μL of total volume in 1.5-mL Eppendorf tubes or in 384-well plates for Malachite green assays. Reactions were assembled on ice from stock solutions within the ranges described in **Table 1**, with most reactions run (unless otherwise noted) with 15 mM HEPES pH 7.3, 60 mM potassium acetate, 4-10 mM magnesium acetate, 30 mM ammonium acetate, 2 mM ATP, 1 mM GTP, 1 mM CTP, 1 mM UTP, 2 mM each of 20 amino acids, 8 mM creatine phosphate, 5 mM DTT, 0.05 mM spermidine, 2% w/v PEG3350, 0.5 U/uL RNase inhibitor (Clontech, Mountain View, CA, USA), 0.1 mg/mL T7 polymerase (made in house following the protocol from Swartz et al., 2004),^78^ 0.33 mg/mL creatine phosphokinase (from rabbit muscle, Sigma-Aldrich), 9 nM plasmid or linear DNA, and 50% v/v S30 extract. Reactions were run overnight at room temperature. Plasmid DNA was prepared using the ZymoPure II Plasmid Midiprep Kit followed by an ethanol precipitation and linear expression templates were prepared by PCR and subsequent cleanup using the Zymo DNA Clean and Concentrator (Zymo Research, Irvine, CA, USA).

The amount of active firefly luciferase produced was quantified by activity test. Four microliters of CFPS sample were added to 30 μL of ONE-Glo Luciferase Assay System (Promega, Madison WI, USA) in a white 96-well plate (Costar 3693, Corning, Corning, NY, USA). Luminescence was measured at 26 °C every 2 minutes for 20 minutes using a Bio Tek (Winooski, VT, USA) Synergy 2 plate reader. For each reaction, the maximum relative light units (RLU) was used to compare to a linear standard curve of recombinant luciferase taken under the same conditions in cell-free buffer added to the ONE-Glo mixture.

### mRNA Detection with Malachite Green Assay

Reactions were assembled in triplicate in 10 μL volumes on ice as described above with 0.02 mM malachite green dye, 30 mM HEPES pH 7.7, 60 mM potassium acetate, 10 mM magnesium acetate, 60 mM ammonium acetate, 2mM ATP, 1 mM GTP, 1 mM CTP, 1 mM UTP, 0.1 mM each of 20 amino acids, 8 mM creatine phosphate, 5 mM DTT, 0.1 mg/mL T7 polymerase (made in house following the protocol from Swartz et al., 2004),^78^ 0.33 mg/mL creatine phosphokinase (from rabbit muscle, Sigma-Aldrich C3755-3.5KU), 3 nM plasmid DNA, or water as a control for background signal and 50% v/v S30 extract. Bacterial reactions were assembled as described in Silverman et al. 2019^61^ with added T7 polymerase.

Kinetic cell-free reactions were assembled on ice in triplicate and pipetted into a 384-well plate (Grenier BioOne 781096) avoiding bubbles. Plates were sealed (Bio-Rad MSB1001) and both sfGFP fluorescence (emission/excitation: 485/528, gain 50) and malachite green fluorescence (emission/excitation: 615/650, gain 100) were measured every 10 minutes overnight for 8 hours at 25 ºC on a BioTek Synergy H1M plate reader.

### Codon Optimization and Plasmid Construction

Codon optimization was conducted on our in-house firefly luciferase sequence by hand based on previous work.^79^ Residues N, D, A, Y, and F were optimized based on reported relative translation efficiencies and all other residues were optimized based on tobacco chloroplast codon usage. Some residues without reported translation rates were left unoptimized to allow for synthesis by Integrated DNA Technologies. Inserts were ordered from Integrated DNA Technologies (Coralville, IA) and cloned into the pJL1 backbone by the method described in Gibson et al.^80^ Inserts included an overhang of anywhere from 27-73 nucleotides with the NdeI and SalI cut sites on pJL1 to facilitate Gibson assembly without the need for primers.

### RBS Library Design and Construction

RBS were designed with the RBS Library Calculator in predict mode with the following sequence (aataattttgtttaactttaagaaggagVBHDHYBD). Of the 4374 variants designed, 104 were selected that sampled the range of the transcription initiation rate of the designs. These were ordered from Twist Bioscience as variations on the pJL1-lucME plasmid with the malachite green aptamer on the 3’ end. Library members were amplified by PCR, purified with a Zymo ZR-96 Clean and Concentrator kit, and quantified with the Promega Quantifluor kit. All DNA was then diluted to an end dilution of 5.63nM and tested as linear expression templates in cell-free reactions in 384-well plates as described above, except that reactions were incubated for 20 hours and optimized cell-free conditions were used for the chloroplast reactions.

## Supporting information

Supplemental RBS data

## Acknowledgments

L.C. thanks Yasushi Yukawa for guidance on chloroplast purification and lysis, Eszter Majer for fruitful discussions, and Holly Ekas for guidance on high throughput DNA quantification and purification for cell-free applications. M.C.J. and C.A.V. were funded by the DARPA 1KM program [HR0011-15-C-0084]. We thank Ashty Karim for his help in editing the manuscript.

