## Supplemental Figures for "A chloroplast cell-free system for measuring ribosome binding site strengths"

**Supplementary Information**

**Supplementary Figures**

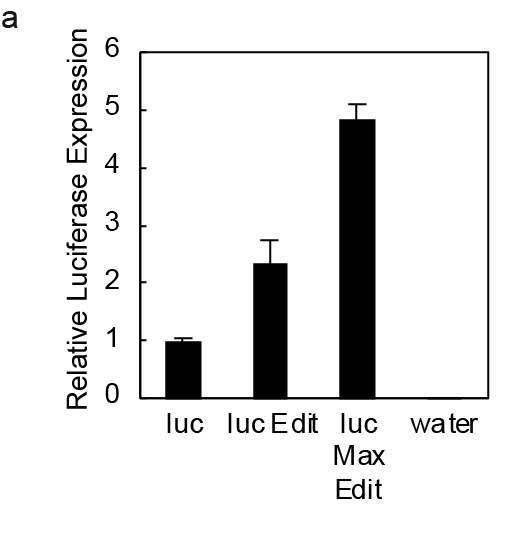

**Figure S1. Codon optimizing our reporter luciferase gene increases CFE.** Three different versions of firefly luciferase were assayed: (1) our standard luciferase used in *E. coli* extracts (Luc), (2) luciferase with N, D, A, Y, and F optimized for the highest rate of translation^105^ (Luc Edit), and (3) luciferase edited for the highest rate of translation and additionally edited to have a higher A/T content to mirror the chloroplast genome (Luc Max Edit, LucME). Experiments were run in triplicate. Values show means with error bars representing standard deviations (s.d.) of three independent experiments (n=3).

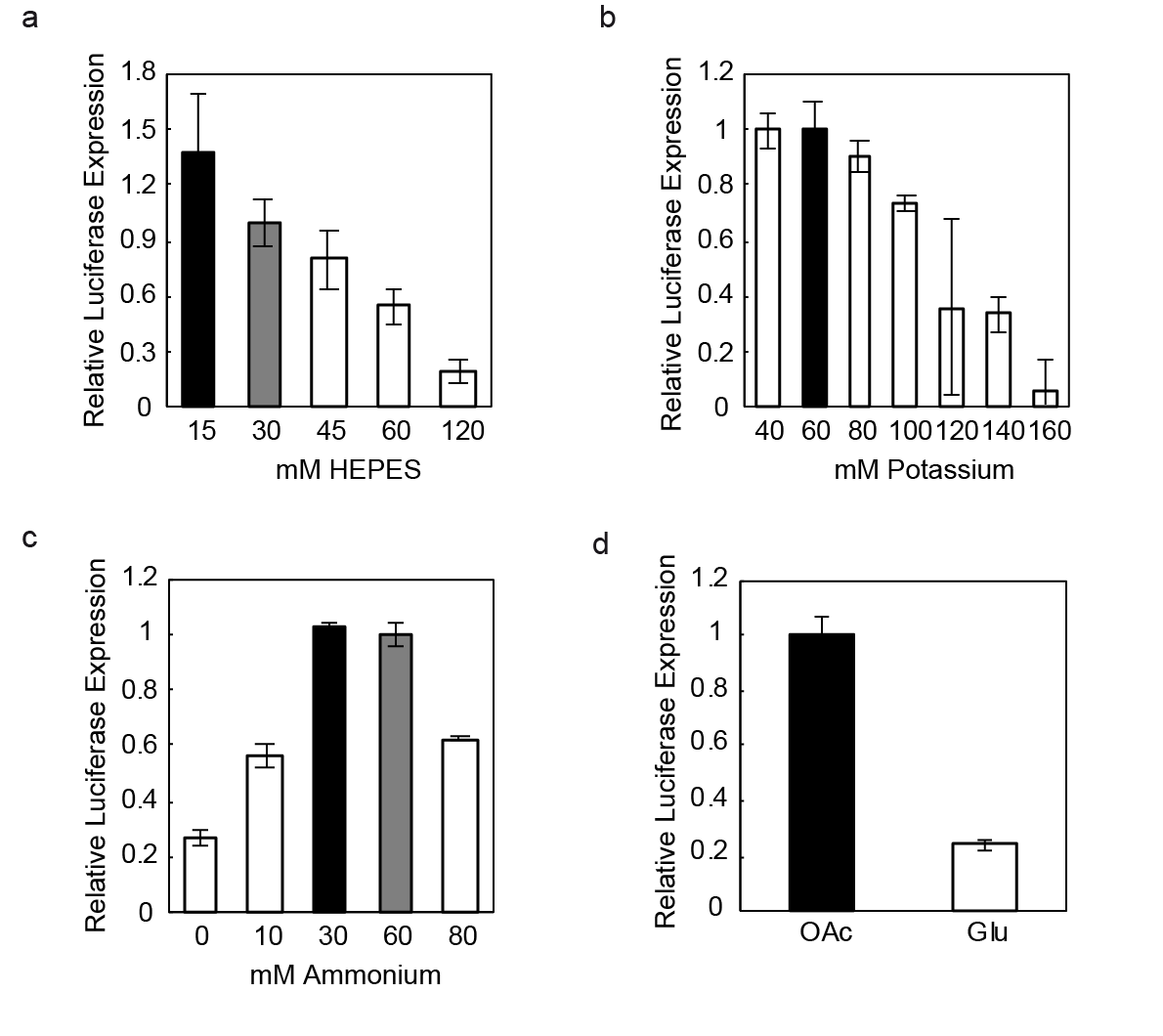

**Figure S2. Dependence of chloroplast-based CFE reactions on different buffer and salt conditions.** (a) HEPES concentration. (b) Potassium acetate concentration. (c) Ammonium concentration. (d) Acetate (OAc) versus glutamate (Glu) salts. Values show means with error bars representing standard deviations (s.d.) of at least three independent experiments (n=3).

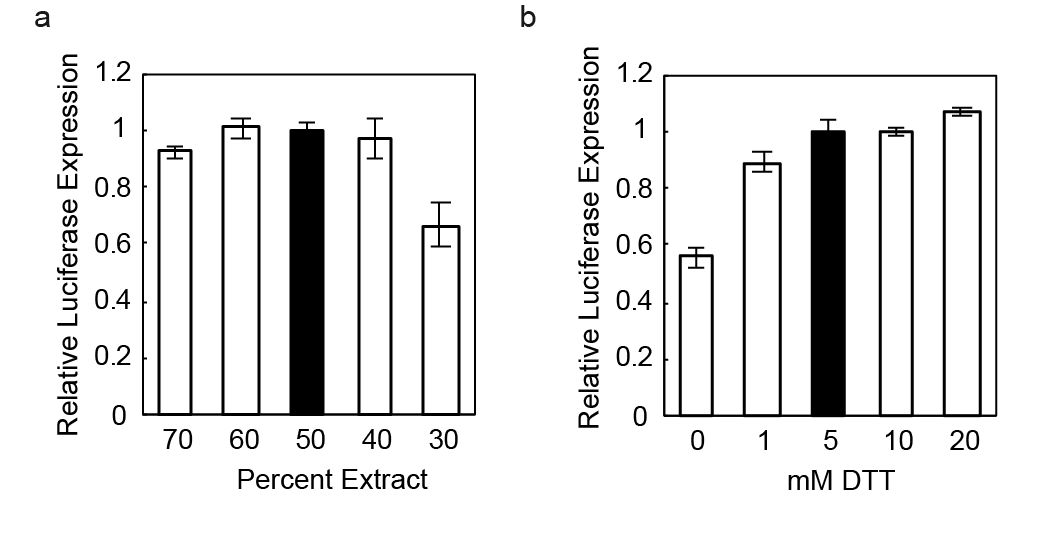

**Figure S3. Dependence of chloroplast-based CFE reactions on percent extract, DTT, and energy substrate.** (A) Extract volume fraction. (B) DTT concentration. Values show means with error bars representing standard deviations (s.d.) of at least three independent experiments.

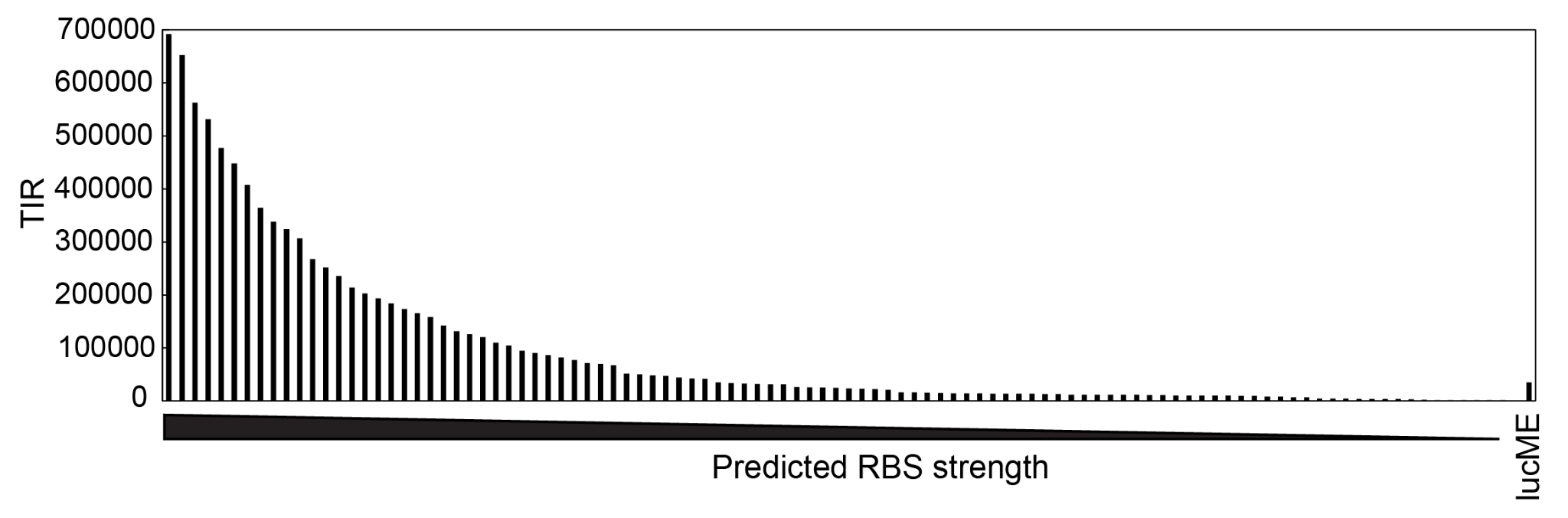

**Figure S4. Predicted RBS strengths of 104-member RBS library.** Predicted translation initiation rate of the RBS designed in the Salis RBS calculator as compared to the control RBS used with the lucME-MGA construct, represented by “lucME”. RBS 1 through 104 are plotted left to right.

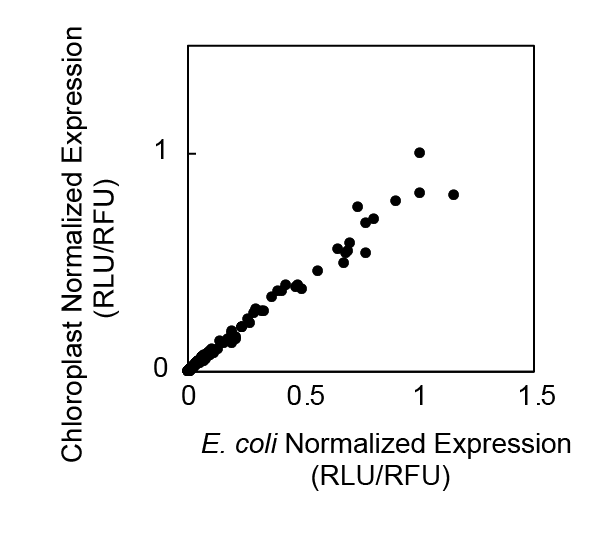

**Figure S5. Normalized gene expression comparison of RBS library in chloroplast versus *E. coli* CFE systems.** Normalized protein expression data (RLU) divided by max transcription value (RFU) for the chloroplast vs *E. coli* system are well correlated (i.e., proteins/transcripts).

**Supplementary Tables**

**Table S1**

|  | **Chloroplast cell-free gene expression**  **(This work: optimized)** | **Chloroplast cell-free gene expression**  **(This work: tested range)** | **Chloroplast cell-free translation only**  **(Yukawa et al., 2007)** | ***E. coli* CFPS**  **(Kwon and Jewett, 2015)** |
| --- | --- | --- | --- | --- |
| **Magnesium** | 4-8mM | 2-14mM | 9mM | 8mM |
| **Amino acids** | 2mM | 0-4mM | 0.04mM | 2mM |
| **Energy regeneration** | CP/CK |  | CP/CK | PEP |
| **Creatine phosphate** | 8mM | n/a | 8mM | n/a |
| **Creatine kinase** | 0.33mg/mL | n/a | 8mg/mL | n/a |
| **DNA** | 9 nM | 1.5-9 nM | 200 fmol/µL mRNA | 13.3 µg/mL |
| **Potassium** | 60mM | 20-200mM | 80mM | 130mM |
| **Ammonium** | 30mM | 0-100mM | 60mM | 10mM |
| **Anion (major species)** | acetate | Acetate, glutamate | Acetate | Glutamate |
| **pH** | 7.3 | n/a | 7.7 | 7.2 |
| **Buffer concentration** | 15mM | 15-120mM | 30mM | 57mM |
| **Extract volume percent** | 0.5 | 30-70% | 10 A280 U/mL | 0.27 |
| **RNase Inhibitor** | 0.5U/µL | 0-1U/µL | 1U/µL in extract before dialysis | n/a |
| **Crowding agent** | PEG3350 | Ficoll 400, PEG600, PEG3350, PEG8000 | n/a | n/a |
| **Crowding agent concentration** | 2% | 0-4% | n/a | n/a |
| **Spermidine** | 0.5mM | 0-1mM | n/a | 1.5mM |
| **DTT** | 5mM | 0-20mM | 5mM | n/a |
| **Nucleotides** | 2mM ATP; 1mM each GTP, CTP, UTP | not tested | 2mM ATP  0.2mM GTP | 1.2mM ATP  0.85mM GTP, CTP, UTP |

**Table S1. Physiochemical optimization ranges of typical cell-free systems.** We conducted physiochemical optimizations in chloroplast extract and present the optimum amount as well as the optimization range of these reaction components as they compare to a previously published translation-only chloroplast system^71^ and the *E. coli* cell free expression system.^88^

**Table S2**

| **RBS** | **Sequence** | **TIR** | **Chloroplast**  **RFU/RLU** |
| --- | --- | --- | --- |
| RBS1 | AATAATTTTGTTTAACTTTAAGAAGGAGGTAATTTGATG | 692190 | 0.47827 |
| RBS2 | AATAATTTTGTTTAACTTTAAGAAGGAGGTAATTGTATG | 652238 | 0.68571 |
| RBS3 | AATAATTTTGTTTAACTTTAAGAAGGAGGTCGTTTAATG | 562525 | 0.76623 |
| RBS4 | AATAATTTTGTTTAACTTTAAGAAGGAGGTCGATTAATG | 531705 | 0.67873 |
| RBS5 | AATAATTTTGTTTAACTTTAAGAAGGAGGTTTATTTATG | 477548 | 1.15253 |
| RBS6 | AATAATTTTGTTTAACTTTAAGAAGGAGGTAGATGTATG | 447883 | 0.49096 |
| RBS7 | AATAATTTTGTTTAACTTTAAGAAGGAGGTATATGAATG | 408062 | 0.46337 |
| RBS8 | AATAATTTTGTTTAACTTTAAGAAGGAGGTTGATGTATG | 364639 | 0.35888 |
| RBS9 | AATAATTTTGTTTAACTTTAAGAAGGAGGTAATTCAATG | 338040 | 0.73258 |
| RBS10 | AATAATTTTGTTTAACTTTAAGAAGGAGGTTGACTAATG | 324315 | 0.64671 |
| RBS11 | AATAATTTTGTTTAACTTTAAGAAGGAGGTATACGTATG | 306643 | 0.76749 |
| RBS12 | AATAATTTTGTTTAACTTTAAGAAGGAGGGATTTTAATG | 267229 | 0.69461 |
| RBS13 | AATAATTTTGTTTAACTTTAAGAAGGAGGGAGATGTATG | 251760 | 0.31754 |
| RBS14 | AATAATTTTGTTTAACTTTAAGAAGGAGGTCGACTTATG | 235410 | 0.80474 |
| RBS15 | AATAATTTTGTTTAACTTTAAGAAGGAGGTCACCGGATG | 213728 | 0.29389 |
| RBS16 | AATAATTTTGTTTAACTTTAAGAAGGAGGGAGATTGATG | 203039 | 0.23259 |
| RBS17 | AATAATTTTGTTTAACTTTAAGAAGGAGGTAAACCTATG | 193450 | 0.89671 |
| RBS18 | AATAATTTTGTTTAACTTTAAGAAGGAGGGAGATTAATG | 183899 | 0.27015 |
| RBS19 | AATAATTTTGTTTAACTTTAAGAAGGAGGTTACTCTATG | 173519 | 0.69887 |
| RBS20 | AATAATTTTGTTTAACTTTAAGAAGGAGGTATACTTATG | 165503 | 1.00555 |
| RBS21 | AATAATTTTGTTTAACTTTAAGAAGGAGGGATTTCAATG | 158369 | 0.39016 |
| RBS22 | AATAATTTTGTTTAACTTTAAGAAGGAGGGTATTCAATG | 142661 | 0.40563 |
| RBS23 | AATAATTTTGTTTAACTTTAAGAAGGAGGGATACCGATG | 131282 | 0.0624 |
| RBS24 | AATAATTTTGTTTAACTTTAAGAAGGAGGGAGACTGATG | 125867 | 0.03608 |
| RBS25 | AATAATTTTGTTTAACTTTAAGAAGGAGGTTTCTGAATG | 120480 | 0.00817 |
| RBS26 | AATAATTTTGTTTAACTTTAAGAAGGAGGGAGACCGATG | 109842 | 0.03982 |
| RBS27 | AATAATTTTGTTTAACTTTAAGAAGGAGGGTACTTGATG | 104579 | 0.19245 |
| RBS28 | AATAATTTTGTTTAACTTTAAGAAGGAGACTATTTGATG | 94827 | 0.12838 |
| RBS29 | AATAATTTTGTTTAACTTTAAGAAGGAGGCAATTTGATG | 90528 | 0.20267 |
| RBS30 | AATAATTTTGTTTAACTTTAAGAAGGAGATTTACGTATG | 86272 | 0.23347 |
| RBS31 | AATAATTTTGTTTAACTTTAAGAAGGAGGGATTTCTATG | 82005 | 0.56171 |
| RBS32 | AATAATTTTGTTTAACTTTAAGAAGGAGATTTATTTATG | 76821 | 0.0208 |
| RBS33 | AATAATTTTGTTTAACTTTAAGAAGGAGGGAGACGGATG | 71690 | 0.03247 |
| RBS34 | AATAATTTTGTTTAACTTTAAGAAGGAGATTTACTAATG | 69749 | 0.2801 |
| RBS35 | AATAATTTTGTTTAACTTTAAGAAGGAGAGTATTCGATG | 67580 | 0.15343 |
| RBS36 | AATAATTTTGTTTAACTTTAAGAAGGAGGCTTATCTATG | 51537 | 0.03607 |
| RBS37 | AATAATTTTGTTTAACTTTAAGAAGGAGAGTTTCGAATG | 50143 | 0.42284 |
| RBS38 | AATAATTTTGTTTAACTTTAAGAAGGAGATAGATCGATG | 48698 | 0.13709 |
| RBS39 | AATAATTTTGTTTAACTTTAAGAAGGAGACCGTTGAATG | 47221 | 0.23537 |
| RBS40 | AATAATTTTGTTTAACTTTAAGAAGGAGGCCGTCTAATG | 44306 | 0.07427 |
| RBS41 | AATAATTTTGTTTAACTTTAAGAAGGAGGTATCTGTATG | 42429 | 0.11322 |
| RBS42 | AATAATTTTGTTTAACTTTAAGAAGGAGACCTACTAATG | 41410 | 0.26033 |
| RBS43 | AATAATTTTGTTTAACTTTAAGAAGGAGCGTATTTAATG | 34756 | 0.17062 |
| RBS44 | AATAATTTTGTTTAACTTTAAGAAGGAGATTTTTCTATG | 33823 | 0.07807 |
| RBS45 | AATAATTTTGTTTAACTTTAAGAAGGAGCGCGTTTTATG | 32463 | 0.5922 |
| RBS46 | AATAATTTTGTTTAACTTTAAGAAGGAGCGTGACTAATG | 32085 | 0.09105 |
| RBS47 | AATAATTTTGTTTAACTTTAAGAAGGAGACTGACGAATG | 31506 | 0.08383 |
| RBS48 | AATAATTTTGTTTAACTTTAAGAAGGAGATTGACGGATG | 31223 | 0.09361 |
| RBS49 | AATAATTTTGTTTAACTTTAAGAAGGAGGGCTCTCGATG | 26251 | 0.09436 |
| RBS50 | AATAATTTTGTTTAACTTTAAGAAGGAGACTTTTTAATG | 25644 | 0.20754 |
| RBS51 | AATAATTTTGTTTAACTTTAAGAAGGAGAGCAATTAATG | 25035 | 0.04758 |
| RBS52 | AATAATTTTGTTTAACTTTAAGAAGGAGCTTATTGGATG | 24412 | 0.02704 |
| RBS53 | AATAATTTTGTTTAACTTTAAGAAGGAGACAACTTAATG | 23545 | 0.05096 |
| RBS54 | AATAATTTTGTTTAACTTTAAGAAGGAGAGCAATTGATG | 22675 | 0.0809 |
| RBS55 | AATAATTTTGTTTAACTTTAAGAAGGAGCTTTACCTATG | 21890 | 0.01514 |
| RBS56 | AATAATTTTGTTTAACTTTAAGAAGGAGATTATCTGATG | 21121 | 0.09003 |
| RBS57 | AATAATTTTGTTTAACTTTAAGAAGGAGCGTGTCGAATG | 16335 | 0.1913 |
| RBS58 | AATAATTTTGTTTAACTTTAAGAAGGAGGCCTTCCTATG | 15769 | 0.06872 |
| RBS59 | AATAATTTTGTTTAACTTTAAGAAGGAGCCAACCGAATG | 15231 | 0.10015 |
| RBS60 | AATAATTTTGTTTAACTTTAAGAAGGAGCGCACTTAATG | 14672 | 0.05546 |
| RBS61 | AATAATTTTGTTTAACTTTAAGAAGGAGATTTCCTAATG | 14132 | 0.06762 |
| RBS62 | AATAATTTTGTTTAACTTTAAGAAGGAGCGCGCTGAATG | 14057 | 0.00477 |
| RBS63 | AATAATTTTGTTTAACTTTAAGAAGGAGCGCACTTTATG | 13997 | 0.04012 |
| RBS64 | AATAATTTTGTTTAACTTTAAGAAGGAGCTCTATTTATG | 13839 | 0.00298 |
| RBS65 | AATAATTTTGTTTAACTTTAAGAAGGAGCGCATTCAATG | 13776 | 0.09231 |
| RBS66 | AATAATTTTGTTTAACTTTAAGAAGGAGGGTGTCTTATG | 13601 | 0.18848 |
| RBS67 | AATAATTTTGTTTAACTTTAAGAAGGAGCTCACTTGATG | 13530 | 0.32542 |
| RBS68 | AATAATTTTGTTTAACTTTAAGAAGGAGCTTATCGTATG | 13300 | 0.05664 |
| RBS69 | AATAATTTTGTTTAACTTTAAGAAGGAGCCTTATGGATG | 13178 | 0.11226 |
| RBS70 | AATAATTTTGTTTAACTTTAAGAAGGAGCGCACCGTATG | 12065 | 0.02691 |
| RBS71 | AATAATTTTGTTTAACTTTAAGAAGGAGCCCATCGTATG | 11917 | 0.05322 |
| RBS72 | AATAATTTTGTTTAACTTTAAGAAGGAGCTCGATCGATG | 11847 | 0.07961 |
| RBS73 | AATAATTTTGTTTAACTTTAAGAAGGAGCCTAACTTATG | 11773 | 0.04522 |
| RBS74 | AATAATTTTGTTTAACTTTAAGAAGGAGCCCTCTGAATG | 11611 | 0.05957 |
| RBS75 | AATAATTTTGTTTAACTTTAAGAAGGAGAGCGCTGAATG | 11531 | 0.03049 |
| RBS76 | AATAATTTTGTTTAACTTTAAGAAGGAGATTTCCTTATG | 11448 | 0.04371 |
| RBS77 | AATAATTTTGTTTAACTTTAAGAAGGAGATAAACGGATG | 11276 | 0.0769 |
| RBS78 | AATAATTTTGTTTAACTTTAAGAAGGAGCTAACTCTATG | 10264 | 0.10012 |
| RBS79 | AATAATTTTGTTTAACTTTAAGAAGGAGCGAGCTGGATG | 10170 | 0.06555 |
| RBS80 | AATAATTTTGTTTAACTTTAAGAAGGAGATCTTTCGATG | 10052 | 0.01319 |
| RBS81 | AATAATTTTGTTTAACTTTAAGAAGGAGCCTACTGTATG | 9927 | 0.06206 |
| RBS82 | AATAATTTTGTTTAACTTTAAGAAGGAGCTCACCCTATG | 9865 | 0.06303 |
| RBS83 | AATAATTTTGTTTAACTTTAAGAAGGAGCGCTACTGATG | 9635 | 0.05506 |
| RBS84 | AATAATTTTGTTTAACTTTAAGAAGGAGCTCACCCGATG | 9398 | 0.05753 |
| RBS85 | AATAATTTTGTTTAACTTTAAGAAGGAGAGCTATCTATG | 8075 | 0.0453 |
| RBS86 | AATAATTTTGTTTAACTTTAAGAAGGAGCTCTACGAATG | 7937 | 0.06235 |
| RBS87 | AATAATTTTGTTTAACTTTAAGAAGGAGCGCTCTTGATG | 6812 | 0.01978 |
| RBS88 | AATAATTTTGTTTAACTTTAAGAAGGAGCCTGTTCTATG | 6620 | 0.02657 |
| RBS89 | AATAATTTTGTTTAACTTTAAGAAGGAGCTCTCTCTATG | 4700 | 0.03568 |
| RBS90 | AATAATTTTGTTTAACTTTAAGAAGGAGCTCTTTCTATG | 4507 | 0.02576 |
| RBS91 | AATAATTTTGTTTAACTTTAAGAAGGAGCGCTCCGGATG | 4213 | 0.00528 |
| RBS92 | AATAATTTTGTTTAACTTTAAGAAGGAGCTCTACCTATG | 3908 | 0.01422 |
| RBS93 | AATAATTTTGTTTAACTTTAAGAAGGAGCTTTCCTGATG | 3814 | 0.03287 |
| RBS94 | AATAATTTTGTTTAACTTTAAGAAGGAGCTCATCCTATG | 3703 | 0.01267 |
| RBS95 | AATAATTTTGTTTAACTTTAAGAAGGAGCGCTTCGTATG | 3567 | 0.01195 |
| RBS96 | AATAATTTTGTTTAACTTTAAGAAGGAGCCATCTTTATG | 3358 | 0.00814 |
| RBS97 | AATAATTTTGTTTAACTTTAAGAAGGAGCCTTCCGAATG | 1798 | 0.00256 |
| RBS98 | AATAATTTTGTTTAACTTTAAGAAGGAGAGCATCTAATG | 1696 | 0.00266 |
| RBS99 | AATAATTTTGTTTAACTTTAAGAAGGAGCGCTTCCGATG | 1612 | 0.00175 |
| RBS100 | AATAATTTTGTTTAACTTTAAGAAGGAGCCTTTCTGATG | 1448 | 0.00464 |
| RBS101 | AATAATTTTGTTTAACTTTAAGAAGGAGCCTTTCTTATG | 1318 | 0.00315 |
| RBS102 | AATAATTTTGTTTAACTTTAAGAAGGAGCTCTTCTGATG | 1271 | 0.00116 |
| RBS103 | AATAATTTTGTTTAACTTTAAGAAGGAGAGCATCTGATG | 1183 | 0.00136 |
| RBS104 | AATAATTTTGTTTAACTTTAAGAAGGAGCTCTTCCTATG | 934 | 0.00071 |
| lucME | AATAATTTTGTTTAACTTTAAGAAGGAGATATACATATG | 35197 | 1.00 |

**Supplementary Table S2. RBS used in this study.** A collection of 104 RBS was designed using the Salis RBS calculator predict mode for *E. coli* to demonstrate modulating translation in chloroplast cell free extracts. Predicted translation initiation rates are listed and RBS were selected semi-randomly from 4374 results to encompass the range of RBS predicted translation initiation rates. RBS were assembled into the pJL1 plasmid backbone by Twist and used as a template for linear expression templates.
